# Modeling of lung-liver interaction during infection in a human microfluidic organ-on-a-chip

**DOI:** 10.1101/2023.06.01.543192

**Authors:** Susanne Reinhold, Christian Herr, Yiwen Yao, Mehdi Pourrostami, Felix Ritzmann, Thorsten Lehr, Dominik Selzer, Yvonne Kohl, Daniela Yildiz, Hortense Slevogt, Christoph Beisswenger, Robert Bals

**Affiliations:** Department of Internal Medicine V – Pulmonology, Allergology and Critical Care Medicine, Saarland University, D-66421 Homburg, Germany; Helmholtz-Institute for Pharmaceutical Research Saarland – HIPS, Molecular Therapies for Lung Disease, D-66123 Saarbrücken, Germany; Clinical Pharmacy, Saarland University, D-66123 Saarbrücken, Germany; Fraunhofer Institute for Biomedical Engineering IBMT, D-66280 Sulzbach, Germany; Experimental and Clinical Pharmacology and Toxicology, Center for Molecular Signaling (PZMS), Saarland University, D-66421 Homburg, Germany; Host Septomics Group, Centre for Innovation Competence (ZIK) Septomics, University Hospital Jena, 07745 Jena, Germany

## Abstract

**Background:** Infections of the respiratory tract such as pneumonia or COVID-19 cause high mortality and morbidity worldwide. Organ-on-a-chip (OC) technologies have been developed in the last years to establish human-based disease models, to study basic disease mechanisms and to provide a tool to speed up drug development. The aim of this study was to establish a lung-liver microfluidic system to study the interaction of both organ modules during infection.

**Methods:** A two organ (lung / liver) microfluidic system was established using primary human bronchial (HBECs) or alveolar type epithelial cells (ATC) for the lung module and Huh-7 cells for the liver module. Inactivated non typeable *Haemophilus influenzae* (NTHi) and *Pseudomonas aeruginosa* PAO1 (PAO1) were applied to the lung module. Secreted mediators were screened by dot-blot analysis and quantified. The effect of lung epithelial bacterial stimulation on the liver cell transcriptome was analyzed by mRNA sequencing.

**Results:** Lung and liver cells established stable cultures in a circulatory microfluidic system. Activation of HBECs or ATCs with NTHi or PAO1 resulted in the secretion of multiple inflammatory mediators into the microfluidic medium including TNF-α, monocyte chemotactic protein-1 (MCP-1) and macrophage inflammatory protein-3 (MIP-3). Addition of lung cells and application of bacterial onto the HBECs module resulted in the gross change of the transcriptome of the liver cell module. Gene ontology enrichment analysis showed the induction of various pathways involved in host defense, metabolisms, repair, and acute phase response.

**Interpretation:** In conclusion, a two-organ lung/liver microfluidic system was established to study the interaction of the organ modules during infection. Mediators released from epithelial culture modules into the microfluidic circulation after exposure to bacterial pathogens significantly modify the gene expression patterns of liver cells.

**Funding:** This research was funded by the German Federal Ministry of Education and Research (BMBF), 031L0153 VISION “Alternativmethoden zum Tierversuch” and the Dr. Rolf M. Schwiete Stiftung.

## Introduction

Infections of the respiratory tract such as pneumonia or COVID-19 cause high mortality and morbidity worldwide. Infections cause local organ damage and in addition systemic effects in the whole organism, which contribute to the outcome of the disease. *In vitro* models have been used to study various aspects of the interaction between microorganisms and lung epithelial cells. Further, the predictive power of animal experiments in drug development has been criticized, and it has been proposed that in several aspects human cell models may reflect the *in vivo* situation better. This has accelerated the model development within the 3R strategy to replace animal experiments. However, models allowing to study the interaction between different organs during infectious diseases are limited.

Organ-on-a-chip (OC) technologies have been developed in the last years with the aims to establish human-based disease models to study basic disease mechanisms and to provide a tool to speed up drug development (1–4). One important goal of OC development is to generate models that help in the development of drugs and evaluate the toxicology of diverse substance such as drugs or environmental substances. Nevertheless, it is difficult to model many aspects of an intact organisms in a complex *in vitro* OC system (3).

Various systems have been developed for studying lung or liver cells in OC systems. Culture of lung cells has been developed in the last decades with several areas of interest. Isolation of stem cells for airway or alveolar cells has been developed and the role of the interaction between epithelial cells and the mesenchymal niche have been highlighted (5, 6). Epithelial cells can be cultured in a variety of approaches comprising conventional submersed, air liquid interface or organoid cultures with or without coculture with supportive cells. Lung OC systems have been developed in a variety of setups including mechanical stretching to model ventilatory movements (7) (“breathing lung-on-a-chip”). A small airway OC system recapitulated COPD-like inflammatory processes (8, 9). Liver cells have been used in various setups for OC systems (10).

The combination of several organ systems in a fluidic system opens the possibility to study inter-organ interactions and thus approaches the aim to study biological processes or the impacts of impact of drugs in organisms-like structures. A few studies used multi-OC included lung and liver modules. The combination of human airway epithelial cells (AECs) in ALI culture and HepaRG^TM^ liver spheroids was used to study the modulate the toxicity of aflatoxin B1 by liver cells (11). A similar approach was used to study aflatoxin B1 detoxification in a liver-lung multi-OC using AT1, AT2, and liver spheroids (12, 13). Other multi-OC setups combined intestinal, liver, skin and kidney cells (12, 14), neurospheres and liver cells (11, 14–16). OC systems have also been applied to study bacterial infectious events (8, 17–20).

Despite the progress in the field, several obstacles slow down the broad application and expansion of OC technologies in basic and translational science (21). To generate fully differentiated tissue equivalents, it is necessary to provide optimal growth conditions, which are difficult to establish in most OC systems. In multi-OC, several parameters need to be considered: individual culture conditions for the individual organ modules, connecting media, microfluidic system, presence or absence of additional cells/tissues such as fibroblasts, immune or endothelial cells. Infection models in OC multi-OC systems have been rarely established and require additional adjustments such as the application route and dosing of the microorganisms. For lung-OC systems, the use of AT1/2 cells is still difficult and limited by organ-typical differentiation.

The aim of the present study was the evaluation of a lung – liver multi-OC system in infection models. We aimed to characterize how gene expression in liver cells is regulated by infected lung epithelial cells. For the lung cells we applied differentiated air liquid interface (ALI) cultures of airway and alveolar human primary cells.

## Materials and Methods

### Cell isolation and culture

Human bronchial epithelial cells (HBEC) were obtained by brush biopsies from patients subjected to bronchoscopy. Isolated cells were expanded in a feeder cell co-culture, supplemented with 10 µM Y-27632 (ROCK inhibitor, Tocris Bioscience, UK) (22). Human alveolar epithelial cells (alveolar type cells, ATC) were isolated from fresh tumor free lung tissue obtained from surgery and subsequently cut to pieces. The obtained tissue was digested using gentleMACS tubes (Miltenyi, Germany) and 5 ml of a digestion solution per tube (8.4 mg collagenase type I (Life Tech, Germany), 0.5 ml dispase (Corning, USA), and 300 µl DNase1 (Roche, Germany) and filled to 5 ml with PBS). The tubes were placed onto the Miltenyi GentleMACS dissociator (Miltenyi, Germany). After incubation for 30 minutes at 37°C, the cells were filtered through a 70 µm MACSsmart cell strainer (Miltenyi, Germany) and washed out by PBS. After centrifugation at 450 x g for 5 minutes at 4°C, the cell pellet was washed with 5-10 ml FACS buffer (1% FBS (Thomas Scientific; USA), 1mM EDTA (Sigma-Aldrich; USA) in PBS), centrifuged again, filtered through 40 µm cell strainer (Fisher Scientific, USA) and counted via a hemocytometer. To select the ATC cells, the mouse monoclonal primary antibody HT2-280 (Terrace; USA) together with the secondary anti-IgM-magnetic beads were used in a Miltenyi LS column (Miltenyi; Germany) according to the manufacturer’s protocol. Isolated cells were seeded in a cell culture flask in airway epithelial cell growth medium (Promocell, Germany).

HBEC or ATC were seeded on 24 well transwell inserts (growth area: 0.33 cm^2^; pore size: 0,4 µm; Corning, USA) apically coated with 0,1 % collagen from rat tail (collagen type I, rat tail; Merck, Germany) at a density of 1.5 × 10^5^ cells / well and incubated at 37°C with 5% CO_2_. To generate air liquid interface cultures, the apical medium of each cell type was removed after 48 hours to expose the cells to air. For the bronchial epithelial cells, the basolateral medium was changed to the differentiation medium DMEM/ F-12 (1:1) supplemented with 2.5 mM L-glutamine (Fisher Scientific, Germany) + 2% Ultroser G (Cytogen, Germany) and 0.2% primocin (InvivoGen, USA). The alveolar cells were grown and differentiated under ALI conditions in AEpiCM-Alveolar Epithelial Cell Medium (ScienCell, USA) with the Epithelial Growth Supplement EpiGCs (ScienCell, USA) and 2% FBS (Thermofisher Scientific, USA). After the airlift, the transepithelial electrical resistance (TEER) was measured by an Millicell-ERS2 volt-ohm Meter (Merck) daily. After reaching a TEER more than 800 Ω x cm^2^, the cultures were used in the lung-liver-microfluidic chip system.

Huh-7 cells were used for the liver compartment. Cells were grown in DMEM / F-12 (1:1) + 2.5 mM L-glutamin + 10% FBS (Thermofisher Scientific, USA) + 1% penicillin / streptomycin (Thermofisher Scientific, USA) and 7,5 × 10^4^ cells were seeded on circular glass plates (diameter 12 mm), which had been put on the bottom of the wells of the plate. After 2 days, the glass slides with the attached Huh-7 cells were transferred to the liver module microfluidic system (Quasi Vivo; Kirkstall, UK)

### Setup of the microfluidic lung-liver-chip system

A microfluidic system Quasi Vivo (Kirkstall, UK) was used for the co-culture/stimulation assays. The system’s setup is shown in Fig.1A. The reservoir was filled with 8 ml of the differentiation media as described above. The alveolar or airway cells on transwells and the Huh-7 cells on glass slides were placed into the chambers (QV600 for the HBEC/ATC and QV500 for the Huh-7 cells). The flow rate of the pump was adjusted to 500 µl / minute and all tubes were flushed with appropriate media to remove air. Three independent microfluidic circuits were run in parallel at 37°C and 5% CO_2_ for 24 or 48 hours.

**Figure 1:**
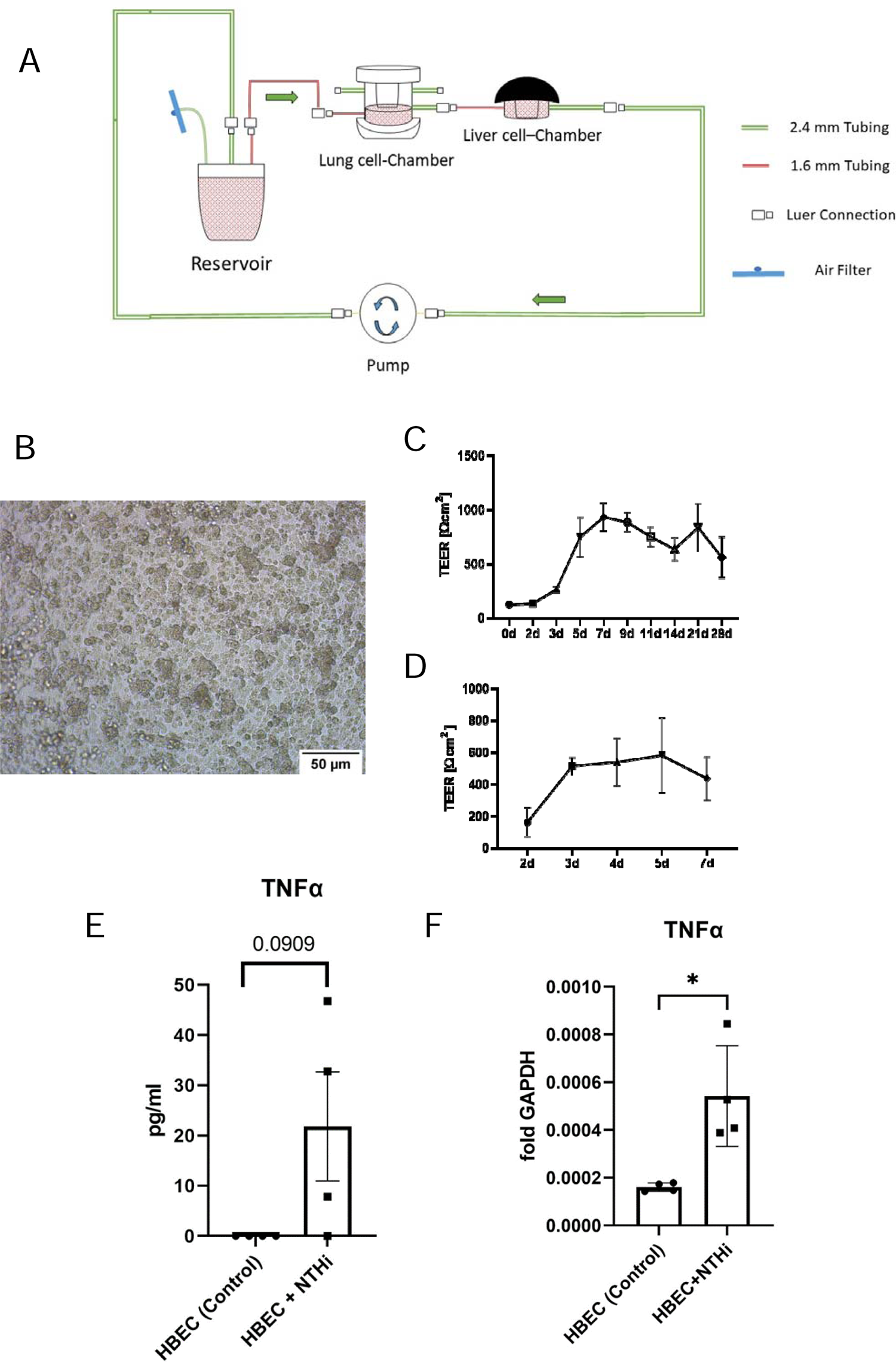
Microfluidic lung-liver microfluidic system. A. Schematic display. The tubing connects the modules lung, liver module and reservoir supported by a peristaltic Parker PF600 pump. The arrows indicate the direction of the liquid flow, the red parts of the tubing have a diameter about 2.4 mm and the green tubing parts of 1.6 mm. B. Micrograph of the confluent HBEC culture on an ALI membrane showing confluent cell growth (bar = 50 µm). C. Development of TEER of HBEC after the removal of the apical medium on day 0 to day 21 cultivation under ALI-conditions. D. Graph of the development of TEER of ATC cultures from the removal of the apical medium on day 0 to day 7 cultivation under ALI-conditions. E, F. Stimulation of HBECs cells with NTHi leads to an increase in the TNF-α concentration (RT-PCR; E) and gene expression (ELISA; F). (unpaired t-test; * P< 0.05, three biological replicates).

### Microbial stimulation models

Heat-inactivated non-typeable *Haemophilus influenzae* (clinical isolates, NTHi) and *Pseudomonas aeruginosa* PAO1 (ATCC 15692, PAO1) were used for the stimulation of the lung cells. Bacteria were stored in frozen glycerol stocks. NTHi was inoculated into 50 ml brain-heart-infusion broth (Roth, Germany) with 1% supplement B (BD Difco™; BD, Germany) and incubated overnight at 37°C. The cultures were collected and centrifuged at 2000 x g for 15 min, the cell pellet was washed with Dulbeccos Phosphate Buffered Saline (DBPS, Sigma-Aldrich, USA). Subsequently, the bacteria were inactivated by heat (70 °C for 15 minutes), cooled on ice and lysed with ultrasound for 45 seconds. The total protein content was analyzed by biochinonic acid-assay (Pierce™ BCA Protein Assay Kit, Thermofisher Scientific, USA). The total protein content of the NTHi lysate was adjusted to 2.5 mg/ml. To validate the inactivity of the bacteria, the NTHi lysates were plated onto chocolate agar. PAO1 was processed similar using LB-Medium (Roth, Germany) at 37°C and the total protein content was adjusted to 1 mg/ml.

For application of the heat-inactivated bacteria, the circuit was stopped and NTHi (20 µl of the bacterial suspension at 50 µg/ml) and PAO1 (20 µl of the bacterial suspension at 20 µg/ml) were applied onto the surface of the HBEC or ATC cultures. For the control group, 20 µl of the appropriate medium was added to the surface. The pump was started again and run for 24 or 48 hours. Then, the medium in the reservoirs was collected and the cells from the chambers were lysed with 350 µl Lysis Buffer RA1 (Macherey-Nagel, Germany) supplemented with 3.5 µl ß-mercaptoethanol (Sigma, USA) for further analysis.

### Characterization of the epithelial secretome by dot-blot analysis and LUMINEX-analysis

To preselect potential cytokines of interest, the expression profile of multiple inflammatory mediators in the media was analyzed by the Proteome Profiler Human XL-Cytokine Array Kit (R&D – Systems, USA) as indicated in the manufacturer’s protocol. The blots were visualized using chemiluminescent detection. The optical densities of the detected analytes were quantified by the software ImageJ 1.53k (National Institute of Health, USA).

The concentrations of selected mediators were quantified by a premixed human multi-analyte Kit (R&D-Systems), containing the analytes angiogenin, MCP-1, MIP-3α, IP-10, IL-1ra, IL-33, BDNF, RANTES, Gro-alpha, IL-1β, IL-12p70 and MIF.

### Quantification of the cytokine concentration after stimulation of HBECs

To quantify the cytokine-concentration secreted by the HBECs after the stimulation with NTHi, we used an enzyme-linked immunosorbant assay (ELISA, Human DuoSet ELISA; R&D Systems) detecting TNF-α according to the manufacturers protocol.

### RT-qPCR and bulk transcriptome analysis of liver and lung cells

To determine the expression of specific genes, RT-qPCR was performed using the SensiMix SYBR&Fluorescein Mix (Meridian; USA) according to the manufacturer’s protocols. Sequences of used primers are: ***CXCL8:*** forward 5’-GTTTTTGAAGAGGGCTGAG-3’ reverse 5’-TTTGCTTGAAGTTTCACTGG-3’; ***SAA1***: forward 5’-CTGCAGAAGTGATCAGCG-3’ reverse 5’-ATTGTGTACCCTCTCCCC-3’; ***TNFA:*** forward 5’-TGCACTTTGGAGTGATCGGC-3’ reverse 5’-ACTCGGGGTTCGAGAAGATG-3’.

To analyze the total transcriptome of the Huh-7-cells in the liver module, RNA was extracted using the NucleoSpin RNA kit (Macherey-Nagel, Germany). The RNA concentration and quality were determined by Nanodrop 8000 spectrophotometer (Thermofisher Scientific, USA). RNA was subjected to bulk sequencing (Nova Seq 6000; PE50) at the Helmholtz Centre for Infection Research (Braunschweig, Germany)

### Limulus amebocyte lysate (LAL)-Endotoxin-Assay

To exclude the presence of bacterial particles in the media, which may have overcome the epithelial barrier and ended up in the media of the circulation system, lipopolysaccharide (LPS) was quantified in the media by a limulus amebocyte lysate assay (Pierce Chromogenic Endotoxin Quant Kit, Thermofisher Scientific, USA) according to the manufacturer’s protocol. We added heat inactivated NTHi (dilution of the stock with the total protein content of 2500 µg/ml was 1: 5 × 10^6^) and PAO1 (dilution of the stock with the total protein content of 1000 µg/ml was 1: 2 × 10^6^) as positive controls. The bacteria were diluted to fit in the detection range of the LAL-assay. Dilutions were made in the respective cellular media as described for NTHi and PAO1.

### Bioinformatics

For analysis of the mRNA sequencing results, all data were aligned with STAR (version: 2.7.3a) with default parameters, count table were generated based on gene ID using featureCounts (version: 2.0.1) with default settings. Alignment and gene counts were generated against the (GRCh38) genome assembly. DESeq2 package (version: 1.36.0) in R (version: 4.2.1) was used to find differentially expressed genes, the threshold for significance was an adjusted P-value < 0.05 and log_2_FoldChange >|1|. Heatmaps were generated using pheatmaps (version: 1.0.12). Functional enrichment analysis, including gene ontology (GO) was carried out using shinyGO web tool.

### Statistical analysis

Data were analyzed by using Graph-Pad Prism 9.0 (for Windows, GraphPad Software, San Diego, California USA). Differences between more than two groups were analyzed using one way ANOVA with Tukeýs post-test. In case of two treatment groups and satisfied parametrical assumptions a Student’s t-test was used. If the parametric assumptions were not reached, Mann-Whitney-U was used and indicated in the figure legends. Results were considered as statistically significant with p < 0.05.

## Results

### Establishment and validation of the lung liver chip

A two-organ microfluidic system was setup using cultures of HBEC or ATC for the lung module and Huh-7 cells for the liver module. Conventional microscopy revealed a closed epithelial layer (Fig. 1B). We measured the transepithelial electrical resistance (TEER) as a sign for the integrity of the epithelium. Fig. 1C,D shows the typical development of TEER measurements for HBEC (C) or ATC (D) cultures, respectively.

### Microbial stimulation of lung cells

In a next step we established microbial exposures to the lung module. We used two clinically relevant bacterial species PAO1 and NTHi. Bacteria were heat-inactivated to avoid overgrowth of the cultures and to allow for long-term analysis. Bacteria were applied to the apical surface of the lung epithelial cultures and the microfluidic system was run for 24 or 48 hours.

To analyze the conditions for microbial stimulation, RT-PCR was used to evaluate the induction of typical host defense genes after the stimulation with NTHi. Compared to the unstimulated control group, the stimulation with NTHi caused an increase in the TNF-α concentration in the supernatant (Fig. 1E). The expression of TNF-α was increased in the NTHi group (Fig. 1F).

### Microbial stimulation of the lung module results in the release of multiple inflammatory mediators

Using the infection models, we first aimed to characterize the secretome of lung epithelial cells as present in the circulating culture medium. 24 and 48 hours after bacterial exposure, media were collected and subjected to dot blot screening. Fig. 2A displays the results after 24 hours of incubation with NTHi and PAO1 of HBEC and Table 1 summarizes the results (24 and 48 h). These data show that bacterial exposure results in release of various mediators into the systemic, microfluidic circulation.

**Figure 2.**
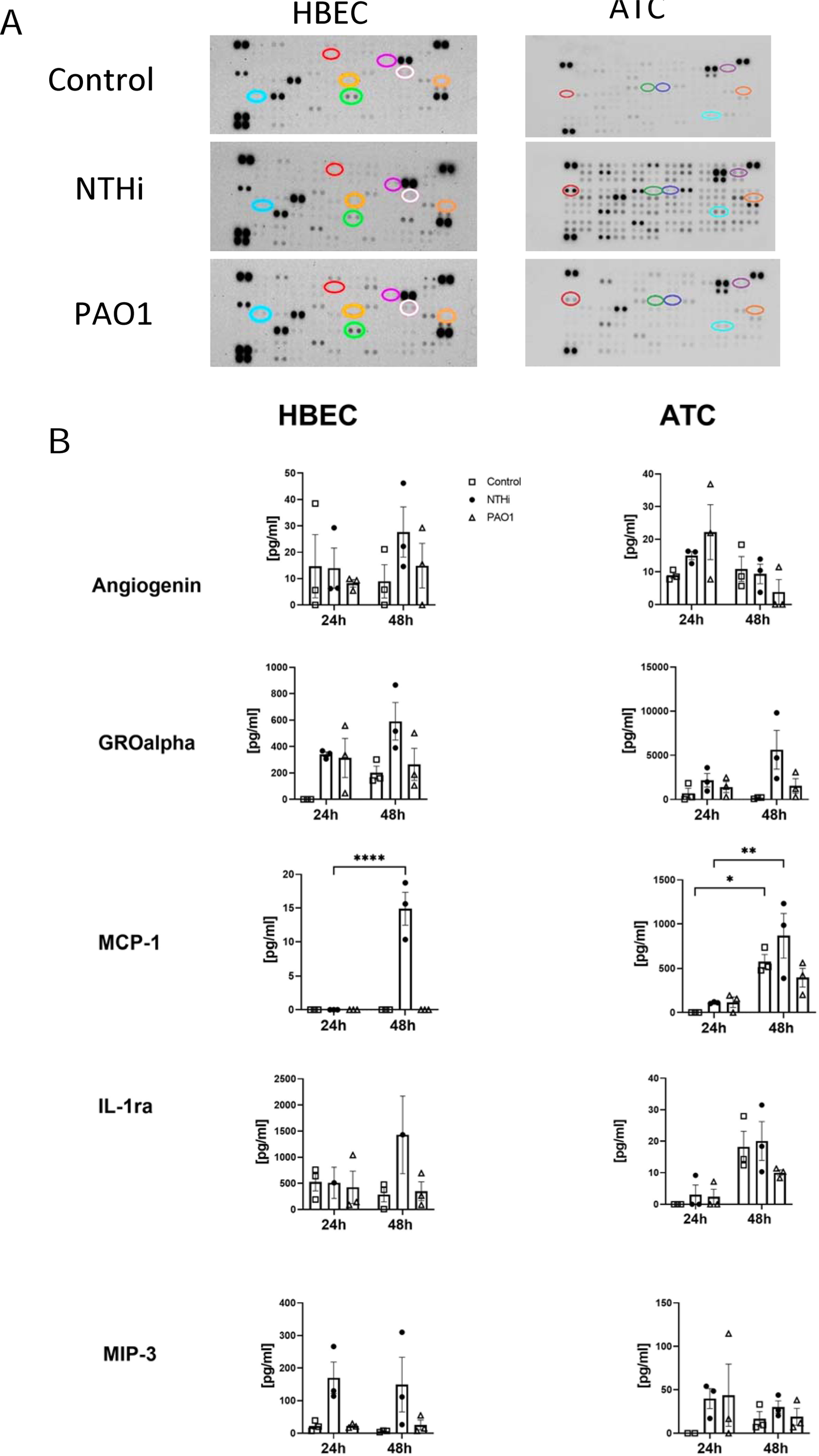
Microbial stimulation of the lung module results in release of mediators in the microfluidic compartment. A. Dot-blot analysis was used for screening. The optical density of the dots indicates the semi-quantitative abundancy of multiple mediators for HBEC and ATC after stimulation with NTHi or PAO1. The most prominent changes were marked with circles in various colours. B. Quantification of selected mediators by Luminex assay. Error bars show the SEM, (n=3), * P<0.05; ** P<0.01; *** P<0.001; **** P<0.0001 after one-way analysis of variance.

**Table 1:** Dot blot analysis of mediator patterns released from HBECs. Optical density of mediators after HBEC-stimulation with NTHi and PAO1 as compared to the control.

|  | 24h |  |  |  | 48h |  |  |  |
| --- | --- | --- | --- | --- | --- | --- | --- | --- |
| | ΔoD NTHi | change | ΔoD PAO1 | change | ΔoD NTHi | change | ΔoD PAO1 | change |
| Adiponectin | 0.0 |  | 666.7 | ↑ | 0.0 |  | -150.0 | ↓ |
| Apolipoprotein A-1 | -56.7 | ↓ | -50.7 | ↓ | 0.0 |  | 0.0 |  |
| Angiogenin | 0.0 |  | -160.0 | ↓ | 300.0 | ↑ | 150.0 | ↑ |
| Angiopoetin-1 | 0.0 |  | -600.0 | ↓ | 0.0 |  | -37.5 | ↓ |
| Angiopoetin-2 | -80.0 | ↓ | -68.6 | ↓ | 0.0 |  | -75.0 | ↓ |
| BDNF | 0.0 |  | -88.9 | ↓ | 0.0 |  | -60.0 | ↓ |
| CD 14 | 0.0 |  | 0.0 |  | -54.5 | ↓ | 0.0 |  |
| CD 30 | -74.7 | ↓ | -97.0 | ↓ | 0.0 |  | -37.5 | ↓ |
| CD 40 Ligand | 0.0 |  | 114.3 | ↑ | 0.0 |  | 0.0 |  |
| Chitinase 3-like 1 | 0.0 |  | -80.0 | ↓ | 0.0 |  | 0.0 |  |
| CFD | 0.0 |  | 0.0 |  | -300.0 | ↓ | -128.6 | ↓ |
| CRP | 0.0 |  | 600.0 | ↑ | 0.0 |  | -900.0 | ↓ |
| Cripto-1 | 0.0 |  | -200.0 | ↓ | 240.0 | ↑ | 0.0 |  |
| Cystatin C | 0.0 |  | -333.3 | ↓ | 0.0 |  | -214.3 | ↓ |
| DPPIV | 0.0 |  | -320.0 | ↓ | 75.0 | ↑ | 0.0 |  |
| EGF | 0.0 |  | -333.3 | ↓ | 0.0 |  | 0.0 |  |
| Emmprin | -40.0 | ↓ | -40.0 | ↓ | 0.0 |  | 0.0 |  |
| ENA-78 | -800.0 | ↓ | -400.0 | ↓ | 0.0 |  | 0.0 |  |
| Endoglin | 0.0 |  | -133.3 | ↓ | 0.0 |  | 0.0 |  |
| Fas Ligand | 0.0 |  | 0.0 |  | 37.5 | ↑ | -37.5 | ↓ |
| FGF basic | 0.0 |  | -114.3 | ↓ | -54.5 | ↓ | -27.3 | ↓ |
| FGF-7 | 0.0 |  | 0.0 |  | -128.6 | ↓ | -128.6 | ↓ |
| FGF-19 | 0.0 |  | -333.3 | ↓ | 0.0 |  | 0.0 |  |
| Flt-3 Ligand | 0.0 |  | 0.0 |  | 150.0 | ↑ | 450.0 | ↑ |
| G-CSF | 36.4 | ↑ | 0.0 |  | 0.0 |  | 60.0 | ↑ |
| GDF-15 | 1.0 | ↑ | 0.0 |  | 0.0 |  | 0.0 |  |
| GM-CSF | -57.1 | ↓ | -85.7 | ↓ | 0.0 |  | 0.0 |  |
| GROalpha | 22.0 | ↑ | -27.8 | ↓ | 1549.2 | ↑ | 0.0 |  |
| Growth Hormone | 0.0 |  | -400.0 | ↓ | 0.0 |  | -150.0 | ↓ |
| HGF | 0.0 |  | 0.0 |  | 75.0 | ↑ | 0.0 |  |
| ICAM-1 | 1400.0 | ↑ | 600.0 | ↑ | 0.0 |  | 0.0 |  |
| IFN-γ | -9.3 | ↓ | -93.0 | ↓ | 0.0 | | -60.0 | ↓ |
| IGFBP-2 | 0.0 |  | 0.0 |  | 60.0 | ↑ | 60.0 | ↑ |
| IGFBP-3 | 0.0 |  | 0.0 |  | -1500.0 | ↓ | 0.0 |  |
| IL-1α | 0.0 | | -53.7 | ↓ | 0.0 | | 600.0 | ↑ |
| IL-1β | 0.0 | | 0.0 | | -37.5 | ↓ | 37.5 | ↑ |
| IL-1ra | 0.0 |  | 40.0 | ↑ | 0.0 |  | -60.0 | ↓ |
| IL-2 | 0.0 |  | -8.3 | ↓ | 0.0 |  | 933.3 | ↑ |
| IL-3 | 0.0 |  | 0.0 |  | -300.0 | ↓ | 0.0 |  |
| IL-4 | -80.0 | ↓ | 0.0 |  | 75.0 | ↑ | 0.0 |  |
| IL-5 | 0.0 |  | -40.0 | ↓ | 0.0 |  | -161.5 | ↓ |
| IL-6 | -96.0 | ↓ | -56.0 | ↓ | 0.0 |  | 0.0 |  |
| IL-8 | 1.8 | ↑ | 1.8 | ↑ | 0.0 |  | 0.0 |  |
| IL-10 | 0.0 |  | 0.0 |  | 0.0 |  | -600.0 | ↓ |
| IL-11 | 0.0 |  | 0.0 |  | 0.0 |  | 450.0 | ↑ |
| IL-12p70 | 0.0 |  | 0.0 |  | -138.5 | ↓ | -207.7 | ↓ |
| IL-13 | 0.0 |  | 0.0 |  | 27.3 | ↑ | -54.5 | ↓ |
| IL-15 | 0.0 |  | 0.0 |  | 0.0 |  | -85.7 | ↓ |
| IL-17A | 0.0 |  | -333.3 | ↓ | 0.0 |  | 0.0 |  |
| IL-18bp-a | 0.0 |  | 0.0 |  | 150.0 | ↑ | 0.0 |  |
| IL-23 | 0.0 |  | 0.0 |  | 0.0 |  | -214.3 | ↓ |
| IL-24 | 109.1 | ↑ | 0.0 |  | 0.0 |  | 0.0 |  |
| IL-27 | 0.0 |  | 0.0 |  | 0.0 |  | -900.0 | ↓ |
| IL-31 | 0.0 |  | 0.0 |  | 0.0 |  | -375.0 | ↓ |
| IL-32 | 0.0 |  | 0.0 |  | 0.0 |  | -180.0 | ↓ |
| IL-33 | 0.0 |  | 0.0 |  | 112.5 | ↑ | 37.5 | ↑ |
| IL-34 | 0.0 |  | 0.0 |  | 54.5 | ↑ | -27.3 | ↓ |
| IP-10 | 0.0 |  | 0.0 |  | 600.0 | ↑ | 900.0 | ↑ |
| I-TAC | 0.0 |  | 0.0 |  | 0.0 |  | -75.0 | ↓ |
| Kallikrein-3 | 0.0 |  | -48.5 | ↓ | 0.0 |  | 0.0 |  |
| Leptin | 0.0 |  | 0.0 |  | -225.0 | ↓ | 0.0 |  |
| LIF | 0.0 |  | 0.0 |  | -150.0 | ↓ | -131.3 | ↓ |
| Lipocalin-2 | 2.1 | ↑ | 3.6 | ↑ | 3.3 | ↑ | 0.0 |  |
| MCP-1 | -31.6 | ↓ | 0.0 |  | 6600.0 | ↑ | 0.0 |  |
| MCP-3 | 0.0 |  | 0.0 |  | 0.0 |  | -900.0 | ↓ |
| M-CSF | 0.0 |  | 0.0 |  | -375.0 | ↓ | -450.0 | ↓ |
| MIF | -93.3 | ↓ | -40.0 | ↓ | 2976.9 | ↑ | -138.5 | ↓ |
| MIG | 0.0 |  | 0.0 |  | -150.0 | ↓ | 750.0 | ↑ |
| MIP-1α/1β | 5000.0 | ↑ | 0.0 | | -150.0 | ↓ | -225.0 | ↓ |
| MIP-3α | -4800.0 | ↓ | 0.0 | | 2228.6 | ↑ | -171.4 | ↓ |
| MIP-3β | 22.2 | ↑ | 0.0 | | 60.0 | ↑ | -150.0 | ↓ |
| MMP-9 | -3.0 | ↓ | -1.8 | ↓ | 0.5 | ↑ | 0.0 |  |
| Myeloperoxidase | 30.8 | ↑ | 15.4 | ↑ | 0.0 |  | 0.0 |  |
| Osteopontin | -57.1 | ↓ | -28.6 | ↓ | -187.5 | ↓ | -131.3 | ↓ |
| PDGF-AA | 52.2 | ↑ | -87.0 | ↓ | 0.0 |  | -144.0 | ↓ |
| PDGF-AB/BB | 0.0 |  | 0.0 |  | -171.4 | ↓ | -214.3 | ↓ |
| Pentraxin-3 | 14.1 | ↑ | -52.5 | ↓ | 0.0 |  | -109.1 | ↓ |
| PF-4 | 0.0 |  | 0.0 |  | -1200.0 | ↓ | -900.0 | ↓ |
| RAGE | 0.0 |  | 0.0 |  | -1200.0 | ↓ | -900.0 | ↓ |
| RANTES | 0.0 |  | 0.0 |  | 0.0 |  | 600.0 | ↑ |
| RBP-4 | -133.3 | ↓ | 0.0 |  | 0.0 |  | 0.0 |  |
| Relaxin-2 | -200.0 | ↓ | 0.0 |  | 0.0 |  | -330.0 | ↓ |
| Resistin | -22.2 | ↓ | -22.2 | ↓ | 0.0 |  | -173.7 | ↓ |
| SDF-1α | 0.0 | | -90.9 | ↓ | 0.0 | | 0.0 | |
| Serpin E1 | -0.5 | ↓ | 0.8 | ↑ | 0.0 |  | 0.0 |  |
| SHBG | -54.5 | ↓ | -66.7 | ↓ | 0.0 |  | -225.0 | ↓ |
| ST2 | 0.0 |  | -88.9 | ↓ | 0.0 |  | -138.5 | ↓ |
| TARC | 0.0 |  | 0.0 |  | -225.0 | ↓ | -300.0 | ↓ |
| TFF3 | 0.0 |  | 0.0 |  | 0.0 |  | 450.0 | ↑ |
| TfR | -76.2 | ↓ | -66.7 | ↓ | -150.0 | ↓ | 0.0 |  |
| TGF-α | 400.0 | ↑ | 0.0 | | 0.0 | | 0.0 | |
| Thrombospondin-1 | 0.0 |  | -83.0 | ↓ | -210.0 | ↓ | 0.0 |  |
| uPAR | 0.0 |  | 0.0 |  | 0.0 |  | -168.8 | ↓ |
| VEGF | -76.9 | ↓ | 0.0 |  | 0.0 |  | -214.3 | ↓ |
| VitaminD-bp | -55.9 | ↓ | -79.6 | ↓ | 0.0 |  | -106.5 | ↓ |
| CD-31 | 0.0 |  | 0.0 |  | -106.5 | ↓ | -116.1 | ↓ |
| TIM-3 | 0.0 |  | 0.0 |  | -207.7 | ↓ | -184.6 | ↓ |
| VCAM-1 | 0.0 |  | 0.0 |  | 0.0 |  | -171.4 | ↓ |

Next, we used ATCs because this cell type is also relevant in infection of the respiratory tract and likely has different properties as compared to airway epithelial cells. ATC cultures were exposed to NTHi and PAO1 and media was analyzed by dot blot analysis (Fig. 2A). Table 2 summarizes the results at 24 hours and 48 hours.

**Table 2:** Dot blot analysis of mediator patterns released from ATCs. Optical density of mediators after stimulation of ATC with NTHi and PAO1 as compared to the control.

|  | 24h |  |  |  | 48h |  |  |  |
| --- | --- | --- | --- | --- | --- | --- | --- | --- |
| | ΔOD<br>NTHi | change | ΔOD<br>PAO1 | change | ΔOD<br>NTHi | change | ΔOD<br>PAO1 | change |
| Adiponectin | 691.4 | ↑ | -61.7 | ↓ | 280.0 | ↑ | 680.0 | ↑ |
| Apolipoprotein A-1 | 1754.9 | ↑ | -9.8 | ↓ | -60.9 | ↓ | -43.5 | ↓ |
| Angiogenin | 1772.7 | ↑ | -90.9 | ↓ | 800.0 | ↑ | 200.0 | ↑ |
| Angiopoetin-1 | 3281.3 | ↑ | 375.0 | ↑ | -40.0 | ↓ | -80.0 | ↓ |
| Angiopoetin-2 | 960.2 | ↑ | 51.1 | ↑ | -25.6 | ↓ | -87.2 | ↓ |
| BAFF | 2346.2 | ↑ | 134.6 | ↑ | -90.9 | ↓ | -72.7 | ↓ |
| BDNF | 2065.6 | ↑ | 204.9 | ↑ | -177.8 | ↓ | -88.9 | ↓ |
| C5/C5a | -5750.0 | ↓ | -500.0 | ↓ | -266.7 | ↓ | 266.7 | ↑ |
| CD 14 | 2843.8 | ↑ | 125.0 | ↑ | 11.8 | ↑ | 117.6 | ↑ |
| CD 30 | 731.7 | ↑ | 146.3 | ↑ | -26.1 | ↓ | 8.7 | ↑ |
| CD 40 Ligand | 940.8 | ↑ | -105.3 | ↓ | 44.4 | ↑ | 66.7 | ↑ |
| Chitinase 3-like 1 | 636.4 | ↑ | 325.2 | ↑ | 12.9 | ↑ | -57.6 | ↓ |
| CFD | 2357.1 | ↑ | -23.8 | ↓ | -114.3 | ↓ | -57.1 | ↓ |
| CRP | 1902.8 | ↑ | 69.4 | ↑ | -93.3 | ↓ | -66.7 | ↓ |
| Cripto-1 | 3500.0 | ↑ | 83.3 | ↑ | -66.7 | ↓ | -133.3 | ↓ |
| Cystatin C | 1555.6 | ↑ | 41.7 | ↑ | -200.0 | ↓ | -66.7 | ↓ |
| DKK-1 | 2180.3 | ↑ | 139.3 | ↑ | -59.3 | ↓ | -81.5 | ↓ |
| DPPIV | 610.5 | ↑ | 232.6 | ↑ | -95.7 | ↓ | 69.6 | ↑ |
| EGF | 0.0 |  | -0.2 | ↓ | -0.5 | ↓ | -0.2 | ↓ |
| Emmprin | 50000.0 | ↑ | 12500.0 | ↑ | 35.3 | ↑ | 35.3 | ↑ |
| ENA-78 | 2309.5 | ↑ | -71.4 | ↓ | 571.4 | ↑ | 57.1 | ↑ |
| Endoglin | 2142.9 | ↑ | -142.9 | ↓ | -85.7 | ↓ | 28.6 | ↑ |
| Fas Ligand | 2875.0 | ↑ | 187.5 | ↑ | 0.0 |  | -200.0 | ↓ |
| FGF basic | 309.8 | ↑ | -18.1 | ↓ | -32.7 | ↓ | -93.9 | ↓ |
| FGF-7 | -1357.1 | ↓ | -571.4 | ↓ | -200.0 | ↓ | 0.0 |  |
| FGF-19 | 1696.1 | ↑ | 68.6 | ↑ | -76.9 | ↓ | -107.7 | ↓ |
| Flt-3 Ligand | -1666.7 | ↓ | -111.1 | ↓ | 800.0 | ↑ | -200.0 | ↓ |
| G-CSF | 4318.2 | ↑ | 363.6 | ↑ | 1200.0 | ↑ | 2400.0 | ↑ |
| GDF-15 | 31.0 | ↑ | 31.3 | ↑ | 15.8 | ↑ | -13.5 | ↓ |
| GM-CSF | -2642.9 | ↓ | -785.7 | ↓ | 80.0 | ↑ | 51.4 | ↑ |
| GROα | 210000.0 | ↑ | 52500.0 | ↑ | 58.1 | ↑ | -23.9 | ↓ |
| Growth Hormone | -1421.1 | ↓ | -52.6 | ↓ | -66.7 | ↓ | 0.0 |  |
| HGF | 4333.3 | ↑ | -333.3 | ↓ | -66.7 | ↓ | 200.0 | ↑ |
| ICAM-1 | 1209.7 | ↑ | -32.3 | ↓ | 61.5 | ↑ | -30.8 | ↓ |
| IFN-γ | 500.0 | ↑ | 5.7 | ↑ | -53.7 | ↓ | -39.0 | ↓ |
| IGFBP-2 | 14000.0 | ↑ | -500.0 |  | 200.0 | ↑ | 0.0 |  |
| IGFBP-3 | 20500.0 | ↑ | -500.0 | ↓ | 0.0 |  | -133.3 | ↓ |
| IL-1α | 831.9 | ↑ | 37.6 | ↑ | -28.6 | ↓ | -97.1 | ↓ |
| IL-1β | 2454.5 | ↑ | -227.3 | ↓ | 0.0 | | -200.0 | ↓ |
| IL-1ra | 2656.3 | ↑ | 62.5 | ↑ | -400.0 | ↓ | 400.0 | ↑ |
| IL-2 | 4666.7 | ↑ | -576.7 | ↓ | -200.0 | ↓ | 125.0 | ↑ |
| IL-3 | 3083.3 | ↑ | 1173.3 | ↑ | -200.0 | ↓ | 125.0 | ↑ |
| IL-4 | 1261.9 | ↑ | -71.4 | ↓ | -1000.0 | ↓ | 0.0 |  |
| IL-5 | 363.8 | ↑ | -46.7 | ↓ | 22.2 | ↑ | -88.9 | ↓ |
| IL-6 | 748.1 | ↑ | 26.7 | ↑ | 44.4 | ↑ | 71.1 | ↑ |
| IL-8 | 2684.2 | ↑ | 2671.1 | ↑ | 0.0 |  | -0.2 | ↓ |
| IL-10 | 1309.5 | ↑ | -47.6 | ↓ | -57.1 | ↓ | 171.4 | ↑ |
| IL-11 | 1253.5 | ↑ | 49.3 | ↑ | -200.0 | ↓ | 40.0 | ↑ |
| IL-12p70 | 880.3 | ↑ | -28.2 | ↓ | -66.7 | ↓ | -133.3 | ↓ |
| IL-13 | 2083.3 | ↑ | -333.3 | ↓ | 133.3 | ↑ | 133.3 | ↑ |
| IL-15 | -10625.0 | ↓ | 0.0 |  | -133.3 | ↓ | -155.6 | ↓ |
| IL-16 | 2937.5 | ↑ | 125.0 | ↑ | -88.9 | ↓ | -44.4 | ↓ |
| IL-17A | 1478.0 | ↑ | 236.3 | ↑ | -59.5 | ↓ | 32.4 | ↑ |
| IL-18bpa | 22000.0 | ↑ | 2500.0 | ↑ | -200.0 | ↓ | -600.0 | ↓ |
| IL-19 | 1038.5 | ↑ | -76.9 | ↓ | -266.7 | ↓ | -200.0 | ↓ |
| IL-22 | 2093.8 | ↑ | 62.5 | ↑ | -160.0 | ↓ | 80.0 | ↑ |
| IL-23 | 4318.2 | ↑ | 227.3 | ↑ | -94.1 | ↓ | -58.8 | ↓ |
| IL-24 | 1805.6 | ↑ | 83.3 | ↑ | 146.7 | ↑ | 53.3 | ↑ |
| IL-27 | 746.9 | ↑ | -30.9 | ↓ | -141.2 | ↓ | -94.1 | ↓ |
| IL-31 | 4833.3 | ↑ | 166.7 | ↑ | 800.0 | ↑ | 0.0 |  |
| IL-32 | 3833.3 | ↑ | 23.8 | ↑ | -142.9 | ↓ | -114.3 | ↓ |
| IL-33 | 5083.3 | ↑ | -250.0 | ↓ | 600.0 | ↑ | -200.0 | ↓ |
| IL-34 | 4916.7 | ↑ | -333.3 | ↓ | -200.0 | ↓ | -266.7 | ↓ |
| IP-10 | -7125.0 | ↓ | -625.0 | ↓ | -200.0 | ↓ | -600.0 | ↓ |
| I-TAC | 1761.9 | ↑ | -47.6 | ↓ | -400.0 | ↓ | 400.0 | ↑ |
| Kallikrein-3 | 1074.5 | ↑ | 114.9 | ↑ | -30.3 | ↓ | -12.1 | ↓ |
| Leptin | 1531.3 | ↑ | -31.3 | ↓ | -40.0 | ↓ | -120.0 | ↓ |
| LIF | 3416.7 | ↑ | 83.3 | ↑ | -257.1 | ↓ | -114.3 | ↓ |
| Lipocalin-2 | 823.0 | ↑ | 373.9 | ↑ | -0.5 | ↓ | -22.4 | ↓ |
| MCP-1 | 1931.4 | ↑ | 186.3 | ↑ | -50.4 | ↓ | -60.7 | ↓ |
| MCP-3 | 3272.7 | ↑ | -45.5 | ↓ | 600.0 | ↑ | 0.0 |  |
| M-CSF | 3403.8 | ↑ | 76.9 | ↑ | 40.0 | ↑ | -240.0 | ↓ |
| MIF | 1500.0 | ↑ | 13.9 | ↑ | -32.7 | ↓ | -106.1 | ↓ |
| MIG | 3818.2 | ↑ | -45.5 | ↓ | 266.7 | ↑ | 66.7 | ↑ |
| MIP-1α/1β | -6375.0 | ↓ | -500.0 | ↓ | -266.7 | ↓ | 0.0 | |
| MIP-3α | 91000.0 | ↑ | 14000.0 | ↑ | -400.0 | ↓ | 600.0 | ↑ |
| MIP-3β | -9125.0 | ↓ | -1125.0 | ↓ | -66.7 | ↓ | -66.7 | ↓ |
| MMP-9 | 1222.2 | ↑ | 152.8 | ↑ | -19.5 | ↓ | -46.6 | ↓ |
| Myeloperoxidase | 409.1 | ↑ | -20.7 | ↓ | 16.0 | ↑ | 0.0 |  |
| Osteopontin | 794.6 | ↑ | -35.7 | ↓ | 22.2 | ↑ | -22.2 | ↓ |
| PDGF-AA | 513.5 | ↑ | 4.5 | ↑ | -51.1 | ↓ | -51.1 | ↓ |
| PDGF-AB/BB | 1416.7 | ↑ | -250.0 | ↓ | -145.5 | ↓ | -109.1 | ↓ |
| Pentraxin-3 | 490.2 | ↑ | 46.3 | ↑ | -42.0 | ↓ | -44.4 | ↓ |
| PF-4 | 6000.0 | ↑ | -250.0 | ↓ | -400.0 | ↓ | -400.0 | ↓ |
| RAGE | 3062.5 | ↑ | -93.8 | ↓ | -200.0 | ↓ | 0.0 |  |
| RANTES | 1833.3 | ↑ | -47.6 | ↓ | -142.9 | ↓ | -57.1 | ↓ |
| RBP-4 | 1365.4 | ↑ | 57.7 | ↑ | 0.0 |  | -200.0 | ↓ |
| Relaxin-2 | 2263.9 | ↑ | 55.6 | ↑ | -600.0 | ↓ | 0.0 |  |
| Resistin | 641.8 | ↑ | 21.3 | ↑ | 44.4 | ↑ | 37.0 | ↑ |
| SDF-1α | 1467.4 | ↑ | 76.1 | ↑ | 155.6 | ↑ | -44.4 | ↓ |
| Serpin E1 | 543.0 | ↑ | 51.1 | ↑ | -0.7 | ↓ | -0.2 | ↓ |
| SHBG | 977.3 | ↑ | 0.0 |  | 114.3 | ↑ | -38.1 | ↓ |
| ST2 | 2071.4 | ↑ | 47.6 | ↑ | -40.0 | ↓ | -40.0 | ↓ |
| TARC | 4090.9 | ↑ | 272.7 | ↑ | 0.0 |  | 0.0 |  |
| TFF3 | 1166.7 | ↑ | -95.2 | ↓ | -142.9 | ↓ | 171.4 | ↑ |
| TfR | 1440.6 | ↑ | 84.2 | ↑ | -22.2 | ↓ | -44.4 | ↓ |
| TGF-α | 1531.3 | ↑ | -187.5 | ↓ | -114.3 | ↓ | -28.6 | ↓ |
| Thrombospondin-1 | 1031.3 | ↑ | 62.5 | ↑ | -15.1 | ↓ | -30.2 | ↓ |
| TNF-α | 1250.0 | ↑ | -57.7 | ↓ | -400.0 | ↓ | -400.0 | ↓ |
| uPAR | 2500.0 | ↑ | 166.7 | ↑ | -66.7 | ↓ | -200.0 | ↓ |
| VEGF | 2125.0 | ↑ | 31.3 | ↑ | 240.0 | ↑ | 40.0 | ↑ |
| VitaminD-bp | 1144.6 | ↑ | 93.4 | ↑ | -21.6 | ↓ | -43.2 | ↓ |
| CD-31 | 709.9 | ↑ | -6.2 | ↓ | -63.2 | ↓ | -94.7 | ↓ |
| TIM-3 | 1319.4 | ↑ | -13.9 | ↓ | -266.7 | ↓ | -66.7 | ↓ |
| VCAM-1 | 611.1 | ↑ | -24.7 | ↓ | -171.4 | ↓ | -85.7 | ↓ |

The dot-blot analysis showed that multiple mediators were found increased in the microfluidic circulation after exposure with microorganisms. The concentrations of selected proteins were quantified using a Luminex assay. The results largely confirmed the data from semi-quantitative dot-blot analysis as displayed in Fig.2B and summarized in Table 3.

**Table 3:** Circulating mediators in the media 24 and 48 hours after bacterial stimulation compared to the unstimulated control group by Luminex-Assay. Data were compared to the control using Student’s T-test: (n=3), * P<0.05; ** P<0.01; *** P<0.001; **** P<0.0001

HBEC
|  | 24h |  |  | 48h |  |  |
| --- | --- | --- | --- | --- | --- | --- |
|  | Control | NTHi | PAO1 | Control | NTHi | PAO1 |
| Angiogenin | 16 ± 23 | 14 ± 14 | 8.6 ± 2.7 | 9.2 ± 11 | 29 ± 17 | 18 ± 10 |
| BDNF | 4.5 ± 7.8 | 15 ± 16 | 6.8 ± 5.9 | 0 ± 0 | 51 ± 34 | 14 ± 12 |
| CCL/RANTES | 1.2 ± 2.1 | 0 ± 0 | 0 ± 0 | 0 ± 0 | 6.3 ± 5.5 | 0 ± 0 |
| CCL2/MCP-1 | 0 ± 0 | 0 ± 0 | 0 ± 0 | 0 ± 0 | **9.8 ± 9.6**<br>** | 0 ± 0 |
| CCL20/MIP-3α | 22 ± 19 | **177 ± 93**<br>* | 23 ± 6.6 | 5.5 ± 4.9 | 161 ± 161 | 32 ± 22 |
| CXCL1/GROα | 712 ± 997 | 2170 ± 1294 | 1469 ± 1143 | 166 ± 90 | 5640 ± 3766 | 1916 ± 1038 |
| CXCL10/IP-10 | 1.3 ± 1.0 | 2.1 ± 1.1 | 0.57 ± 0.49 | 0 ± 0 | 4 ± 4.6 | 0.8 ± 0.26 |
| IL-1β/IL-1F2 | 0 ± 0 | 0 ± 0 | 0 ± 0 | 0 ± 0 | 1.8 ± 3.2 | 0 ± 0 |
| IL-12 p70 | 0 ± 0 | 0 ± 0 | 0 ± 0 | 0 ± 0 | 0 ± 0 | 0 ± 0 |
| IL-1ra/IL-1F3 | 576 ± 336 | 516 ± 516 | 422 ± 532 | 302 ± 261 | 1481 ± 1252 | 386 ± 306 |
| IL-33 | 0 ± 0 | 0 ± 0 | 0 ± 0 | 0 ± 0 | 1.2 ± 2.0 | 0 ± 0 |
| MIF | 2103 ± 2052 | 1739 ± 1110 | 2381 ± 3473 | 630 ± 574 | 4178 ± 3072 | 1014 ± 617 |

ATC
|  | 24h |  |  | 48h |  |  |
| --- | --- | --- | --- | --- | --- | --- |
|  | Control | NTHI | PAO1 | Control | NTHI | PAO1 |
| Angiogenin | 9.6 ± 2.1 | 15 ± 2.1 | 22 ± 14 | 11 ± 6.6 | 9.3 ± 5.2 | 3.8 ± 6.6 |
| BDNF | 0 ± 0 | 0 ± 0 | 0 ± 0 | 0 ± 0 | 0 ± 0 | 0 ± 0 |
| CCL/RANTES | 0 ± 0 | 0 ± 0 | 1.2 ± 2.1 | 1.7 ± 2.9 | 0 ± 0 | 0 ± 0 |
| CCL2/MCP-1 | 0 ± 0 | **112 ± 9.7**<br>**** | 115 ± 101 | 574 ± 132 | 848 ± 409 | 401 ± 191 |
| CCL20/MIP-3α | 0 ± 0 | 40 ± 19 | 43 ± 60 | 20 ± 20 | 30 ± 12 | 18 ± 16 |
| CXCL1/GROα | 0 ± 0 | **343 ± 25**<br>**** | 315 ± 246 | 201 ± 88 | 573 ± 216 | 268 ± 214 |
| CXCL10/IP-10 | 0 ± 0 | 0 ± 0 | 0.13 ± 0.23 | 0.33 ± 0.58 | 0 ± 0 | 0 ± 0 |
| IL-1β/IL-1F2 | 0 ± 0 | 0 ± 0 | 0 ± 0 | 2.5 ± 4.3 | 0 ± 0 | 0 ± 0 |
| IL-12p70 | 0 ± 0 | 0 ± 0 | 16 ± 28 | 11 ± 18 | 0 ± 0 | 0 ± 0 |
| IL-1ra/IL-1F3 | 0 ± 0 | 6.1 ± 11 | 4.8 ± 8.3 | 20 ± 12 | 20 ± 10 | 9.6 ± 1.2 |
| IL-33 | 0 ± 0 | 0 ± 0 | 1.2 ± 2.0 | 2 ± 3.5 | 0 ± 0 | 0 ± 0 |
| MIF | 0 ± 0 | 0 ± 0 | 94 ± 96 | 453 ± 233 | 505 ± 239 | 222 ± 39 |

We also measured whether LPS as prototypical bacterial pattern molecule can be detected in the microfluidic circulation by LAL-assay to rule out that microbial patterns directly stimulate hepatocytes. HBEC and ATC were exposed to NTHi or PAO1 for 24 or 48h and LAL-measurements revealed very low LPS concentration in the microfluidic medium equivalent to dilutions of the bacterial stock solution of 1: 5 × 10^6^ for NTHi or 1: 2 × 10^6^ for PAO1, respectively.

### Microbial stimulation of the lung module drives expression of inflammatory and repair genes in the liver cells

We next investigated whether the release of epithelial mediators into the systemic circulation results in changes of gene expression in liver cells. The lung-liver-chip system was used as described above and the HBECs were stimulated with heat-inactivated NTHI or PAO1. As further control, the epithelial cell module was omitted to characterize the impact of the native epithelial module onto liver cell gene expression. After 24 h RNA was extracted from the liver cells and the transcriptome was analyzed. The liver cell transcriptomes of the following groups were compared:

1. native liver cells (LI) vs. liver cells / lung cells (LILU)
2. liver cells / lung cells (unstimulated) (LILU) vs. liver cells / lung cells (stimulated with NTHi (LILU_NTHi)
3. liver cells / lung cells (unstimulated) (LILU) vs. liver cells / lung cells (stimulated with PAO1 (LILU_PAO1)

We first analyzed the impact of native lung epithelium on liver cell gene expression (LI vs LILU) and found significant alterations in gene expression. A heat map shows the segregation of the experimental groups (Fig. 3A) and a volcano blot visualizes differentially expressed genes (Fig. 3B).

**Figure 3:**
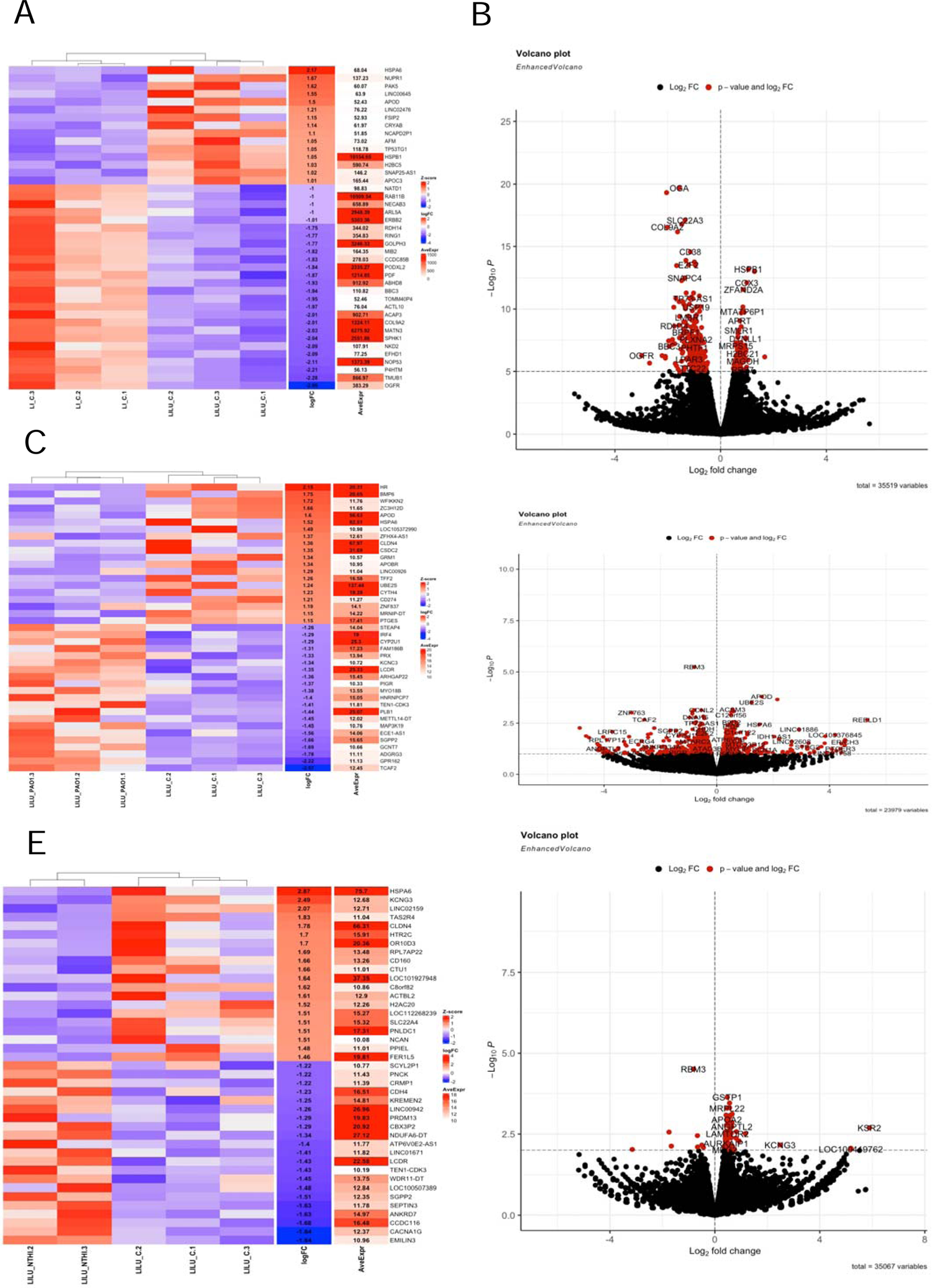
Microbial stimulation of the lung module results in changes of liver cell transcriptome. A-B. The presence of unstimulated HBECs in the microfluidic system result in changes of gene expression in liver cells (LI vs. LILU) displayed by a heatmap (A) showing 20 upregulated and 20 downregulated significant genes out of 345 significantly different genes of Huh-7 cells compared to the connection of Huh-7 and unstimulated HBE cells and a volcano plot showing significant genes in red and non-significant in black (B). C-F. Exposure of the lung module to NTHi (C, D; LILU vs. LILU_HNTi) or PAO1 (E, F; LILU vs. LILU_PAO1) resulted in significant alteration of the liver cell transcriptome as displayed by heatmaps (C, E) or volcano blots (D, F).

We then analyzed the effect of microbial stimulation of the lung module on liver cell gene expression. The application of both bacterial species resulted in modulation of the hepatocyte transcriptome (LILU vs LILU_NTHi; LILU vs. LILI_PAO1). Fig. 3 A, C and E displays the heatmap segregation and Fig. 3 B, D and F differential expression volcano blots after stimulation with PAO1 or NTHi, respectively. The results indicate that addition of the lung module results in significant changes of the liver cell transcriptome.

To evaluate which cellular processes are activated in liver cells, we performed gene ontology enrichment analysis, which showed activation of multiple cellular processes (Fig. 4).

**Figure 4:**
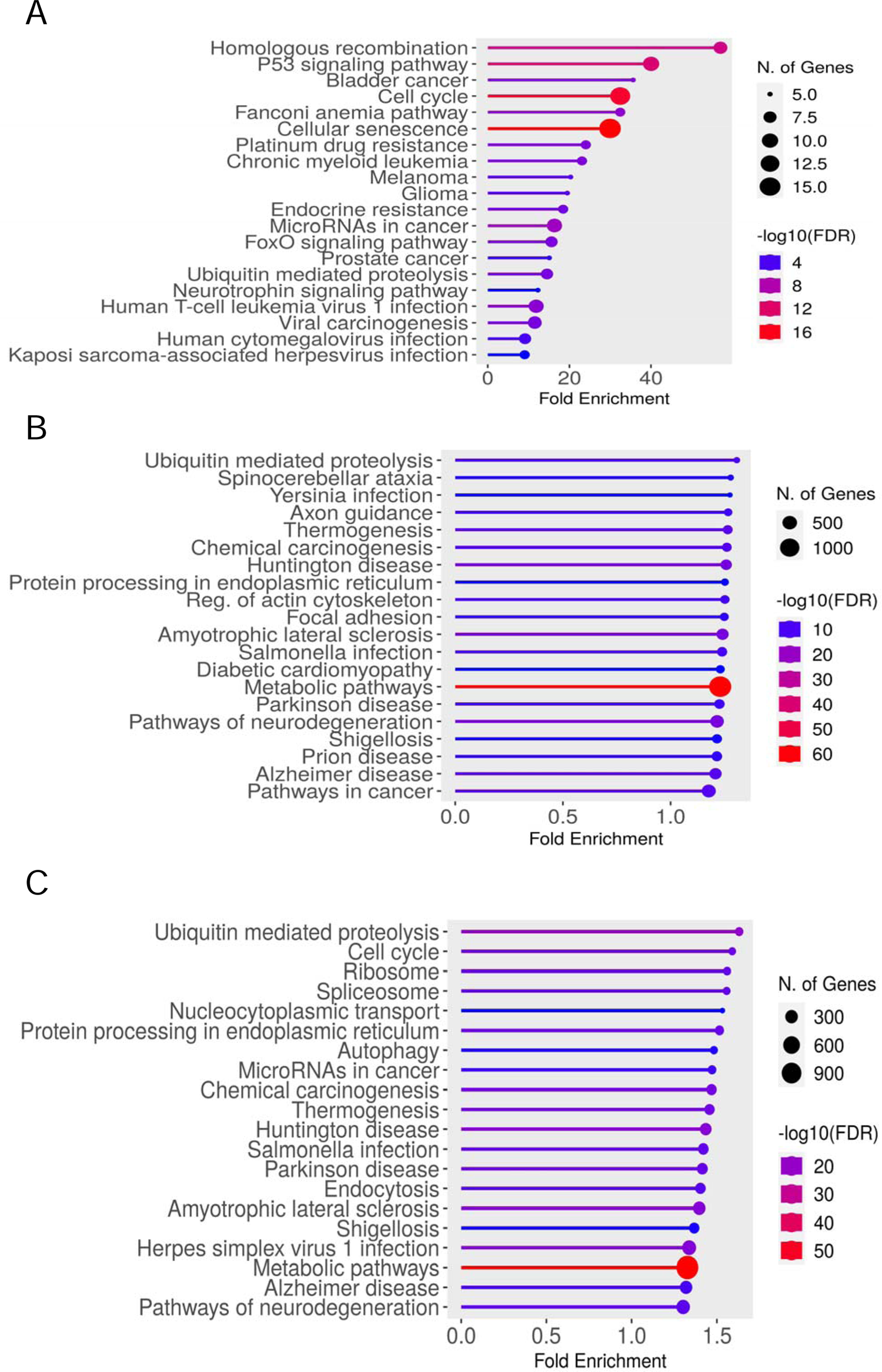
Gene Ontology enrichment shows activation of various signaling pathways. In the microfluidic system. A. Induction of pathways in liver cells (LI) vs. liver cells and unstimulated lung cells (LILU). The DEG of this group of comparison were not enriched for inflammation pathway, host defense pathways and acute phase pathway, only DNA repair pathway is enriched with 172 genes with fold enrichment of 1.41 and FDR cutoff value of 0.05. B and C. Induction of pathways in liver cells with lung cells stimulated with NTHi (LILU_NTHi, B) or PAO1 (LILU_PAO1, C) vs. liver cells and unstimulated lung cells (LILU).

To evaluate if microbial patterns could directly stimulate liver cells, we tested whether heat-inactivated bacteria could activate Huh-7 cells and found that NTHi at the concentration used in the experiments had no detectable effect on the gene expression of typical acute phase proteins of the liver (Fig. 5 A, B). The direct stimulation of Huh-7 with inflammatory cytokines (1000 ng/ml IL-6 and 200 ng/ml TNF-α und IL-1ß) resulted in a significant increase in the expression of acute-phase-proteins (Fig. 5 C, D)

**Figure 5:**
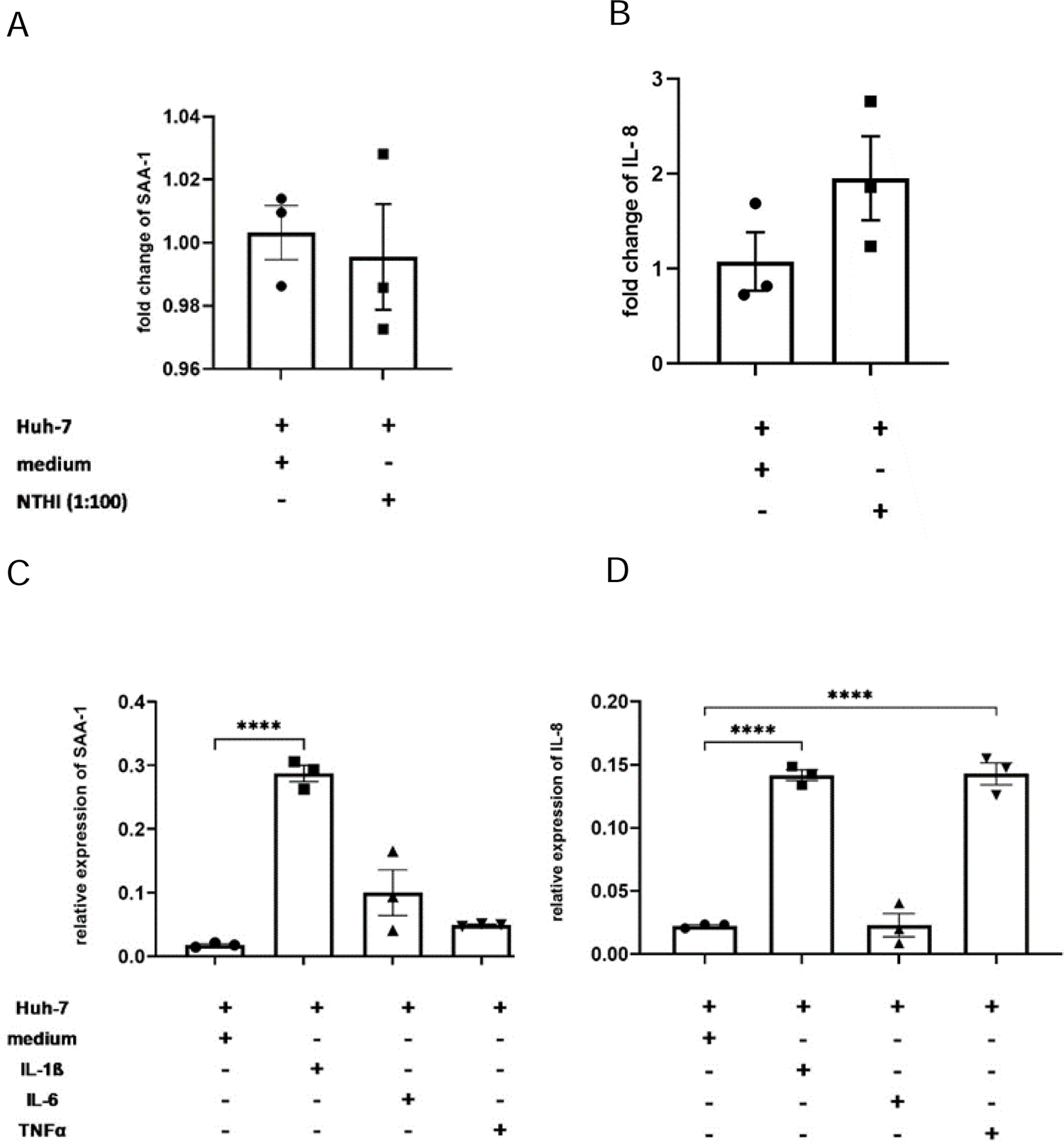
Inflammatory response of Huh-7 to exposure with NTHi and inflammatory cytokines. Exposure of HBECs with NTHi has no significant effect on the gene expression of SAA-1 and IL-8 (A), but inflammatory cytokines significantly induced the gene expression (unpaired Student’s t-test; * P<0.05; ** P<0.01; *** P<0.001; **** P<0.0001, (n = 3).

These data show that mediators released from epithelial culture modules into the microfluidic circulation significantly modify the gene expression patterns of liver cells.

## Discussion

The main finding of this study was that lung epithelial and liver cell cultures can be established in an OC-multi-organ model and that exposure of lung epithelial cells with bacteria activates hepatocytes via secreted mediators. This is the first study that characterized transcriptome regulation in an inter-organ infection model.

Lung OC systems have applied in different setups to model lung tissue and respiratory diseases including ventilatory movements (7, 16) (“breathing lung-on-a-chip”). A small airway OC system recapitulated COPD-like inflammatory processes (8). Lung and liver modules have been combined to study the toxicity of aflatoxin B1 by liver cells (11, 12). In the microfluidic setup used in the present study, we hypothesize that microbial stimulation of the lung epithelial layer results in the secretion of inflammatory mediators into the circulating fluid. These mediators then stimulate liver cells resulting in the acute-phase transcriptomes.

Application of NTHi or PAO to the lung epithelial cells resulted in the secretion of multiple mediators into the microfluidic medium as determined by dot-blot screening and subsequent quantification. The released mediators comprised of MIP-3 and MCP-1 belonging to typical epithelial acute phase proteins. We applied airway epithelial cells (HBEC) and alveolar cells (ATC) as both cell types are involved in host defense and likely react to microbes differentially (23, 24). Indeed, the patterns of released mediators were significantly different between the two cell types, highlighting the need to include HBEC and ATC into experimentation to obtain a comprehensive view on lung host defense.

The main focus of this work was to study the interaction of the lung and liver modules. The sole combination of liver cells with native epithelial cells (HBECs) already caused significant modification of the liver cell transcriptome. Gene ontology analysis showed the induction of multiple pathways, including cell cycle regulation, p53 and other pathways.

Application of NTHi and PAO1 to the epithelial module caused a significant change of the transcriptome of the liver cells with activation of multiple genes. Of note, the expression patterns differed significantly between the two bacterial species, indicating the capability of epithelial cells to discriminate between pathogens.

We investigated whether liver cells could be directly stimulated by microbial patters applied to the apical side of the lung epithelia and potentially translocate into the microfluidic medium. We used LPS as a marker substance as both bacteria applied were Gram negative and found that only traces of this molecule were found in the medium. Stimulation of hepatocytes with bacterial suspensions did not result in the induction of acute phase proteins. It is therefore unlikely that bacterial patterns directly stimulate hepatocytes and changes in the hepatocyte transcriptome are likely induced by epithelial mediators released into the micro-fluidic medium.

This study has limitations and strength. We did not use additional cell types such as fibroblasts or endothelial cells. It is known that fibroblasts help in the differentiation of epithelial cells of the lung or the liver (25). Endothelial cells are important for the establishment of the vascular barrier (26). We decided not to include these cell types to avoid additional complexity in the establishment of the culture conditions. The setup of the multi-organ OC system and the infection protocols comprised multiple parameters such as culture condition, cellular source, microfluidic medium, perfusion parameters and others (27). While we investigated many variables as described in the results section, we did not systematically evaluate all parameters as this would have resulted in an exceeding large number of experiments. We focused on the establishment of critical variables for cell culture, media and dose of bacterial stimulation. We applied heat-inactivated bacterial to better control stimulation conditions and to avoid bacterial overgrowth. In general, the complexity of such multi-organ OC systems challenges the comparability of various published setups. In the present study we used primary lung epithelial cells to avoid biases caused by cell lines. As the focus of this work was on the role of lung epithelial cells, we used human airway and alveolar cells. A liver cell line was used as the secondary module. The use of primary cells allows to use such OC systems to study patient specific properties such as genetic or epigenetic composition of the cells.

In conclusion, we established a lung-liver OC system to study the response to infection. HBEC and ATC stimulated with typical respiratory pathogens released multiple mediators into the microfluidic medium dependent on the cell type and microbial species resulting in distinct patterns of gene induction in hepatocytes.

## Acknowledgments

The authors thank Anja Honecker, Andreas Kamyschnikow, Martina Seibert, and Victoria Weinhold for their excellent support within the areas of laboratory techniques and clinical data management.

